# Lineage-aware evolutionary analysis of hepatitis C virus within-host dynamics

**DOI:** 10.1101/2024.10.15.617766

**Authors:** Lele Zhao, Matthew Hall, Prahalad Giridhar, Mahan Ghafari, Steven Kemp, Haiting Chai, Paul Klenerman, Eleanor Barnes, M. Azim Ansari, Katrina Lythgoe

## Abstract

Analysis of viral genetic data has previously revealed distinct within-host population structures in both untreated and interferon-treated chronic hepatitis C virus (HCV) infections. While multiple subpopulations persisted during the infection, each subpopulation was observed only intermittently. However, it was unknown whether similar patterns were also present after Direct Acting Antiviral (DAA) treatment, where viral populations were often assumed to go through narrow bottlenecks. Here we tested for the maintenance of population structure after DAA treatment failure. We analysed whole-genome next-generation sequencing data generated from a randomised study using DAAs (the BOSON study). We focused on samples collected from patients (N=84) who did not achieve sustained virological response (i.e. treatment failure) and had sequenced virus from multiple timepoints. For each individual, we tracked concordance in nucleotide variant frequencies through time. Using a sliding window approach, we applied sequenced-based and tree-based clustering algorithms across the entire HCV genome. Finally, we reconstructed viral haplotypes and estimated lineage specific within-host divergence rates from the haplotype phylogenies. Distinct viral subpopulations were maintained among a high proportion of individuals post DAA treatment failure. Using maximum likelihood modelling and model comparison, we found an overdispersion of viral evolutionary rates among individuals, and significant differences in evolutionary rates between lineages within individuals. These results suggest the virus is compartmentalised within individuals, with the varying evolutionary rates due to different viral replication rates or different selection pressures. We propose lineage awareness in future analyses of HCV evolution and infections to avoid conflating patterns from distinct lineages, and to recognise the likely existence of unsampled subpopulations.

## Introduction

Infections caused by the hepatitis C virus (HCV) are prevalent globally. While around 30% of infections clear spontaneously, the remaining become chronic if left untreated. Up to 30% of untreated chronic infections result in cirrhosis and liver cancer, leading to an increased risk of death, significantly damaging the health of the individual, and burdening health systems (WHO 2024). In recent years, a range of highly effective Direct Acting Antivirals (DAAs) has proven effective, with efficacy rates up to 95% in some genotypes (Sulkowski et al. 2014; Feld et al. 2015; Wyles et al. 2015; Kowdley Kris V. et al., 2014), although access to diagnosis and treatment remains low in many low and middle income countries (WHO 2024). Without an effective vaccine, reinfections are common in highly affected areas and among at-risk populations (Dietz and Lohmann 2023).

Similar to other RNA viruses, HCV exhibits complex within-host dynamics. It often maintains a unique population structure, where multiple distinct viral populations coexist but are only observed intermittently across sampling time points (Raghwani et al. 2016; Raghwani et al. 2019). Within-host phylogenies have demonstrated the structured viral population of untreated and interferon-treated chronically-infected HCV patients (Gray et al. 2012; Raghwani et al. 2016, 2019; Ramachandran et al. 2011; Riaz et al. 2022). Many biological mechanisms could support such population structure, including physical barriers such as cirrhotic liver tissues (Sorbo et al. 2019; Pérez et al. 2017), or the infection of different cell types (Ducoulombier et al. 2004; Gismondi et al. 2013). HCV also has a very low rate of recombination which acts to prevent the genetic mixing of different subpopulations (Raghwani et al. 2019), whilst compartmentalisation would prevent cells from being coinfected with different subpopulations, reducing the effective rate of recombination still further. Segregating populations could lead to differences in replication rates, generation times and/or rates of evolution.

Understanding viral disease dynamics is an indispensable part of the road toward the World Health Organisation’s 2030 target of eliminating viral hepatitis. The within-host population structure does not only affect drug resistance mechanisms but could also complicate studies looking at disease transmission, because genetically different populations could be circulating between the time of transmission and the time of sampling (Raghwani et al. 2019). This phenomenon will inevitably interfere with molecular epidemiology inferences and surveillance efforts, which usually rely on consensus genetic data.

Studying subpopulations and compartmentalisation of HCV is not straightforward because most viral genetic data is from blood samples, so subpopulations can be mixed. With longitudinal sampling, it has been found that sometimes a single population circulates the body, while at other times, multiple distinct populations coexist in the blood (Raghwani et al. 2016, 2019; Ramachandran et al. 2011). While compartmentalisation has been proposed as the mechanism underlying these dynamics, it is very difficult to investigate because it requires the isolation of viruses infecting the, yet unknown, compartments. These compartments could be different sections of the same liver or various candidate cell types spread across the body.

Without coordinated proactive measures, the high success rates of DAAs may be heavily challenged by emerging viral drug resistance in the future. Moderate to high prevalence of baseline resistance-associated substitutions (RASs) to DAAs have been found in publicly available sequences and treatment studies, occasionally coupled with lowered sustained virological response (SVR) rates (Childs et al. 2019; Gupta et al. 2019; D. Smith et al. 2019; Chen et al. 2016; Palladino et al. 2020; Patiño-Galindo et al. 2016; Takeda et al. 2017). Studies have found that the genetic barriers to DAAs are low across viral genotypes (Kliemann et al. 2016; Patiño-Galindo et al. 2016). RASs that conferred cross-resistance to multiple DAAs sharing the same target were readily selected *in vitro* (Fernandez-Antunez et al. 2023). In addition, the detected RASs were naturally inherent to specific viral genotypes (Vo-Quang et al. 2023; D. Smith et al. 2019; Patiño-Galindo et al. 2016). Different genotypes and demographics vary in their overall SVR rates with DAA treatment (Childs et al. 2019; Dietz et al. 2021; Vo- Quang et al. 2023; Gupta et al. 2019). Detailed characterisations of RASs, especially in light of within-host population structure, are essential for treatment guidelines in resource limited regions as it will help to avoid an increase in drug resistance prevalence and to achieve high levels of SVR post treatment.

An important observation has been the maintenance of viral subpopulations after liver transplantation and treatment with ribavirin and interferon, with distinct subpopulations present prior to transplantation or treatment re-emerging months or years later (Gray et al. 2012; Raghwani et al. 2016). Given the rapid and mass rollout of DAAs, which are highly effective at treating most HCV infections, a key question is whether distinct subpopulations are also maintained in the minority of individuals who fail treatment. To answer this question we analysed short-read whole-genome deep-sequencing data from samples longitudinally collected from patients that failed DAA treatment. We used a variety of analytical methods to identify subpopulations, and if they were present to determine their rates of evolution. As a result of the findings, we suggest the within-host evolutionary analyses of HCV viral populations should be lineage-aware as standard.

## Methods

### Samples, sequencing and bioinformatics

Blood samples were collected from participating patients of the BOSON study (Foster et al. 2015) and permission was granted for the use of samples for further studies. All participating patients provided written informed consent before undertaking any study-related procedures. The BOSON study protocol was approved by each recruiting institution’s review board or ethics committee before study initiation. For the patients that did not achieve sustained viral response (SVR) during the study, samples were collected before, during and post treatment. For several individuals, baseline and post treatment samples were also available as they underwent re- treatment.

RNA was extracted from 500 μl of plasma using the NucliSENS easyMAG system (bioMérieux) into 30 μl of water, of which 5 μl was processed with the NEBNext Ultra Directional RNA Library Prep Kit for Illumina (New England Biolabs) with published modifications to the manufacturer’s protocol (Batty et al. 2013). A 500-ng aliquot of the pooled library was enriched using the xGen Lockdown protocol (Rapid Protocol for DNA Probe Hybridization and Target Capture Using an Illumina TruSeq Library [v1.0]; Integrated DNA Technologies) with a comprehensive panel of HCV-specific, 120-nucleotide DNA oligonucleotide probes (IDT), designed using a published algorithm (Bonsall et al. 2015). The enriched library was sequenced on the Illumina MiSeq v2 platform to produce paired 150-bp reads.

The reads were demultiplexed. Low-quality reads were trimmed using QUASR (v7.0120), and adapters were removed using Cutadapt (v1.7.1). Human reads were removed using Bowtie (v2.2.4). HCV reads were extracted by mapping to 162 ICTV reference sequences (tblastn) for *de novo* assembly with Vicuna (v1.3) and read mapping with *shiver* (Wymant et al. 2018). HCV genotypes were assigned by similarity to ICTV reference sequences. Consensus sequences were generated using *shiver* (v1.7.3) and used to construct a phylogenetic tree. Samples were removed from further analyses if they did not cluster with the other consensus sequences from the same individual. All phylogenetic tree reconstructions in this study were done using IQ- TREE2 with the GTR+F+R6 model (Minh et al. 2020). A flowchart illustrates the sample processing and filtering for the study (Supplementary Figure S1).

### Baseline variant and resistance-associated variant (RAV) frequency trajectories

A total of 266 samples were mapped by *shiver*, which generated base frequency files (i.e. files containing base counts per genomic position) for tracking baseline variant frequencies for 84 longitudinally sampled individuals. A variant position required a minimum depth of 100 reads and a variant allele frequency of at least 10% for at least one time point, to be included in the analysis. After quality filtering and sample availability selection, the variant frequency trajectories were produced for 50 individuals with three or more samples. Variants that were fixed (increased to and remained at above 90%) or purged (decreased to and remained at below 10%) after the first time point were not assessed. For 9 out of the 50 individuals, frequencies of pairs of variants that are closer than 150bp were also extracted from the same sequencing reads to confirm linkage between these variants.

In addition, a list of 65 candidate resistance-associated variants (RAVs) was compiled from previous publications (D. Smith et al. 2019; Ansari et al. 2019, 2017; Wing et al. 2019) including DAA resistant genotypes, DAA resistant genotypes associated with HLA and INFL4 genotypes (Supplementary table 1). Amino acid frequencies of RAVs in individuals of this study were directly extracted from mapped reads. No minimum frequency threshold was applied for the RAVs, but frequency was only assessed when the RAVs were present on two or more mapped reads. All genomic positions are in reference to the H77 genome (GenBank accession: NC_038882).

### Haplotype reconstruction

We used CliqueSNV (Knyazev et al. 2021) to reconstruct haplotypes for the samples in our study (command in Supplementary Text File). One phylogenetic tree per individual was constructed using the haplotypes (aligned with MAFFT (default settings)).

### Population structure in sliding windows

The bam files (N=266) of individuals with two or more longitudinal samples with HCV reads mapped were processed using phyloscanner (phyloscanner_make_trees.py) (Wymant & Hall, et al. 2018). Phyloscanner generated alignments and trees for 210bp sliding windows with 50bp increments across the entire HCV genome. The window width was chosen through visual inspection of analysis results from a random subset of the samples to strike a balance between maximum window width (or read length) and read abundance spanning the window. Next, the k parameter in phyloscanner, determining the routines for identifying contaminant reads and multiple infections was optimised for HCV as described in Supplementary Text File using sequencing reads from 570 patients from the BOSON study. The phyloscanner_analyse_trees.Rscript was used to remove possible contaminants within each sample.

After decontamination, fastbaps (Tonkin-Hill et al. 2019), a hierarchical bayesian clustering tool, was applied to divide the sequences into 2 clusters within each phyloscanner window (Dirichlet prior type = ‘optimise.baps’, number of initial clusters = 2). For each individual patient, genomic windows containing a minimum of 2 sequences from at least 2 different samples were assessed. If both clusters contained sequences from the baseline (pre-treatment) sample and sequences from one or more later (post-treatment) samples, the window was deemed to have sequence-based support for a baseline structured population being maintained throughout treatment. To validate the sequence-based support for the maintenance of population structure, Simmonds’ Association Index (SAI) (Wang et al. 2001), a tree-topology-based statistic for compartmentalisation, was also calculated for the phylogeny constructed for each window. This step took the clusters assigned by fastbaps and tested how stable the clusters were by bootstrapping the phylogenetic trees. A SAI <0.1 (Lewis et al. 2013) indicates strong support for stable clustering.

### Subpopulation divergence rate comparison

To estimate the neutral rate of evolution, the number of synonymous substitutions was counted for each non-baseline haplotype to its closest baseline haplotype. A maximum likelihood model with a Poisson distribution with one overall rate for all individuals in the study was fitted to the observed synonymous substitutions. To accommodate different rates of evolution among individuals, a second maximum likelihood model with a negative binomial distribution was also fitted. Fitting to a negative binomial distribution relaxed the assumption that rates are identical amongst individuals, instead taking each to be drawn from a gamma distribution.

To determine if evolutionary rates vary among subpopulations within the same individuals, we estimated an evolutionary rate per subpopulation. For each individual’s haplotype tree, an exhaustive model comparison analysis was carried out to find the best model of evolutionary rates that explained the distribution of synonymous substitutions across time. Specifically, the non-baseline haplotypes were divided into different hypothetical subpopulations if their closest baseline haplotype and sampling time were different. Linear mixed-effect models were compared for all possible combinations of the hypothetical subpopulations and the best model was determined by the lowest BIC score (see Supplementary Figure S2 for example). The sets of subpopulations and their rates were extracted and categorised into individuals with only one subpopulation, individuals with multiple subpopulations where some subpopulations are missing haplotypes from intermediate time points between the first and last time point of that lineage, and subpopulations that were not missing haplotypes from intermediate time points between the first and last time point of that lineage.

## Results

### Subpopulation frequencies fluctuate across timepoints

Since HCV has been reported to have low rates of recombination (Raghwani et al. 2019), we hypothesised that if different subpopulations have genetic variants unique to them, then the dynamics of subpopulations could be observed by tracking frequency changes of within-host variants through time. We first generated frequency trajectories of genomic variants that were below 50% at Baseline, above 50% at Post-Treatment 12 weeks (PT12) and below 50% at Post-Treatment 24 weeks PT24 (top panel, Figure 1). The majority of these variants were synonymous changes. Similar frequency trajectories were also shown for pairs of variants that are closer than 150bp on the genome (Supplementary Figure S3), further confirming that these frequency concordant variants were on the same genomic background. We then examined the genomic location of these variants (bottom panel, Figure 1). The variant distribution across the HCV genome did not show a specific genomic region responsible for segregating the subpopulations, and also supported the lack of recombination.

**Figure 1.**
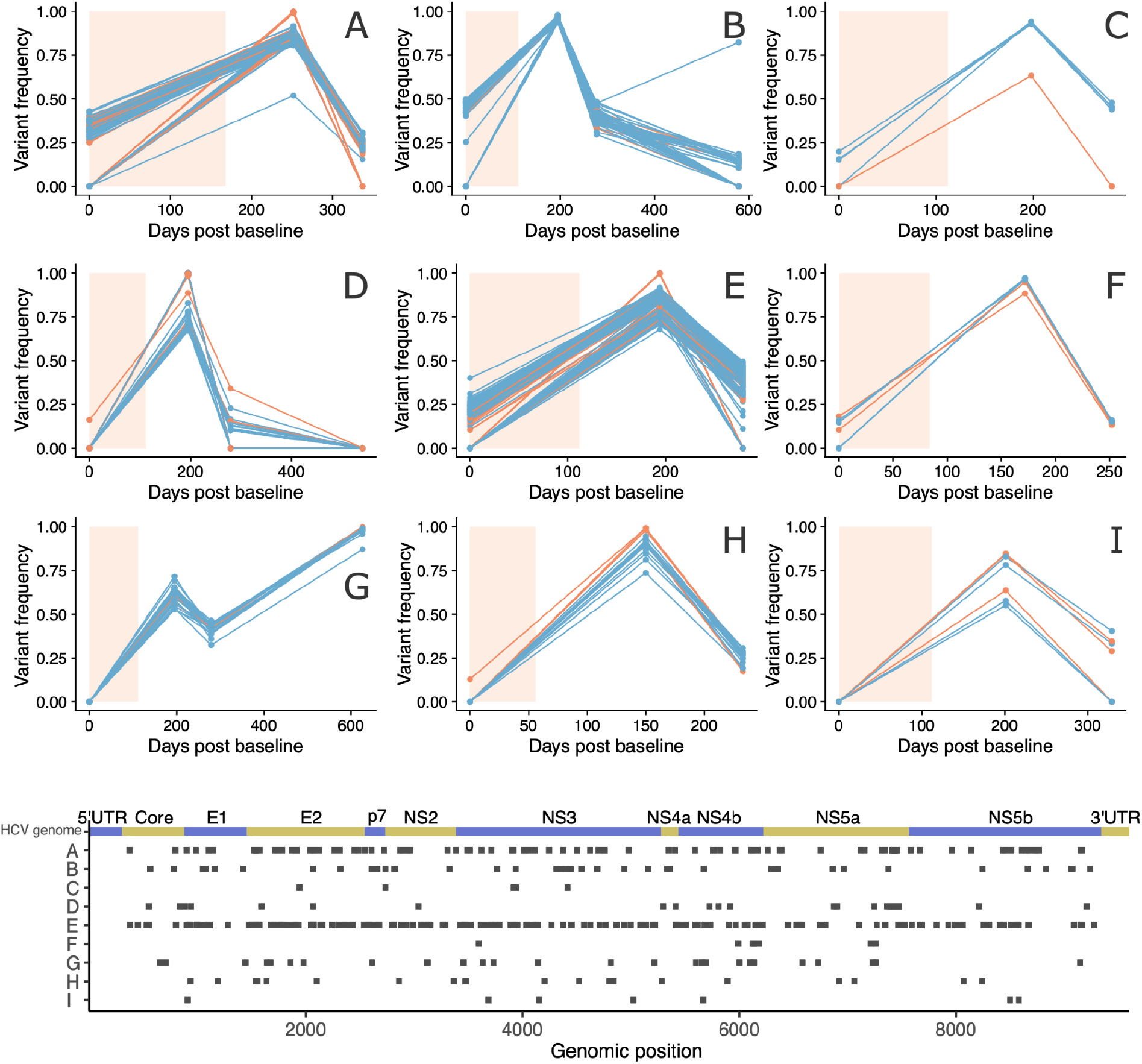
Frequency trajectories of variants that fluctuated (Top) and distribution of genomic sites (Bottom) (patients A-I). The inclusion criterion was variants that changed from below 50% at Baseline (i.e. 0 days post baseline), to above 50% at PT12 (12 weeks post treatment), and decreased to below 50% at PT24 (24 weeks post treatment). Patient B,D and G have a fourth sample after PT24. Synonymous changes are in blue, nonsynonymous changes are in orange and non-coding sites are in grey. The light peach colour shaded region indicates the duration of DAA treatment.

Next, we generated frequency trajectories for all fluctuating variants present in the individuals (Supplementary Figure S4). We observed highly concordant variant frequencies through time, highly suggestive of genetically distinct subpopulations changing in frequency, with different populations represented by unique haplotypes. To explore this further, we reconstructed viral haplotypes for each sample using cliqueSNV, from which we generated haplotype phylogenies for each infection using IQ-TREE2 [see Methods]. The variant trajectories were reflected in many individuals’ haplotype trees (Figure 2). For example, in Figure 2, patient J, the low frequency lineage was present at around 20% at Baseline, dropped to 0% at 12 weeks post treatment (PT12) and reappeared at around 20% at 24 weeks post treatment (PT24). This low frequency lineage was represented by one baseline haplotype and one PT24 haplotype, with both being on the same branch of the haplotype tree. Similar consistencies of haplotype phylogenies with variant trajectories can be seen in other patients, with examples for patients A and B given in figure 2.

**Figure 2.**
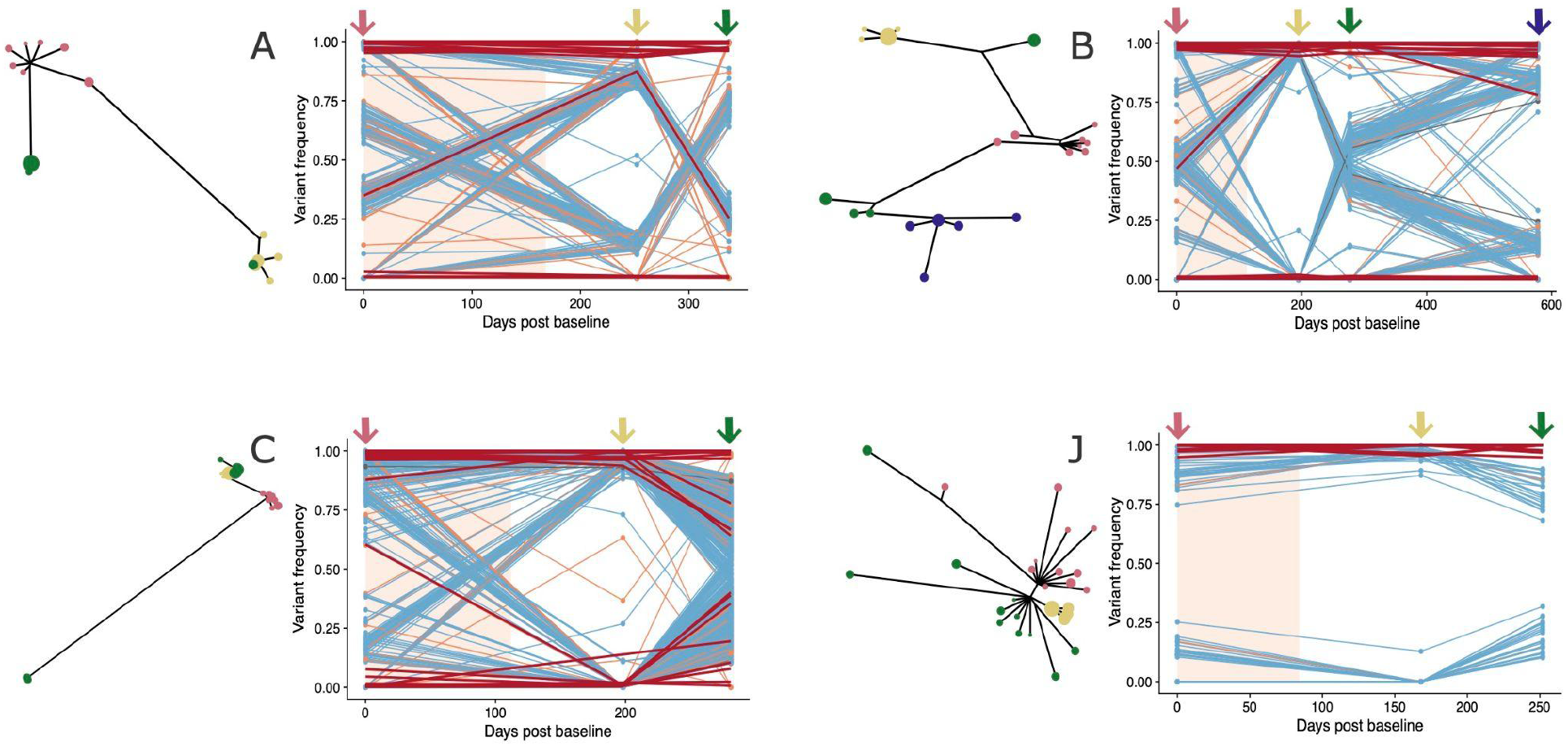
Haplotype phylogenies and variant frequency trajectories across sampling time points (patient A, B, C and J). All nucleotide variants with a frequency above 10% in at least one sample were traced across all sampling time points, with synonymous changes in blue, nonsynonymous changes in orange and non-coding in grey. All resistance-associated variants (RAV) trajectories are in red. The light peach colour shade indicates the duration of the DAA treatment. The coloured arrows on top of the trajectories and the coloured tips in the haplotype phylogenies represent the different sampling timepoints (light red: baseline; yellow: PT12 (12 weeks post treatment); green: PT24 (24 weeks post treatment); purple: BRT (baseline before re-treatment))

However, the correspondence between the allele trajectories and the haplotype trees was not always clear cut, which in some cases may be due to the presence of subpopulations not observed at baseline. For example, in Figure 2: patient C, we speculate the long branch leading to the single haplotype at PT24, was a resurrection of a lineage that was present before, but not at, baseline.

To visualise if the subpopulations exclusively harboured RASs, we plotted RAV frequency trajectories together with the nucleotide variant frequency trajectories (Figure 2 and Supplementary Figure S4). In most of the individuals, there was either a consistent presence (frequency above 90%) or absence (frequency below 10%) of most RAVs throughout the sampling period. A median of 24 RAVs [range 5-28, mean 23.34] were present at equal to or above 50% frequency throughout the sampling period for the 50 assessed individuals. A median of 3 RAVs [range 0-14, mean 4.22] were present below 50% frequency throughout the sampling period for the assessed individuals. For some positions in 20 individuals, we observed RAV frequency fluctuations (i.e. frequency changing from below 50% to above 50% or vice versa) suggesting complex resistance mechanisms. Specifically, NS5B: 90A, 150V, 66T, 120R, 180Q and NS3 67V fluctuated in frequency in 4 individuals, NS5B 90A fluctuated in 5 individuals. In several patients, we observed dynamics suggesting a RAV may have been under selection as a result of the treatment, with a minor subpopulation containing the RAV at baseline becoming the major subpopulation after treatment (Figure 2, patient A-B). However, given the highly dynamic nature of within-host HCV subpopulations and the lack of recombination, we cannot say whether the subpopulation became dominant because it harboured the RAV.

### Pretreatment population structure was retained post treatment

Although the variant frequency trajectories and haplotype phylogenies were suggestive of highly structured within-host populations, we are essentially assuming that variants of similar frequencies are linked (on the same genome). However, because we are using short-read sequencing data we cannot *a priori* make this assumption since most mutations will be on different reads. To further demonstrate the existence of highly dynamic subpopulations, we tested for population structure using sequencing reads spanning sliding windows along the genome, meaning linkage between variants within a given window is known rather than inferred. The short reads meant low resolution in any given window, however, by scanning the entire genome and considering a large number of windows, we were able to build up an aggregate picture for each individual. We used two cross-validating summary statistics to demonstrate the existence of within-host population structure across the entire genome of HCV.

For each window, we first used a Bayesian sequence-based approach (fastbaps) to determine if sequences showed evidence of population structure persisting through the infection. This population structure was then cross-validated using tree topology (SAI). The two summary statistics are very conservative in nature, and therefore a positive result is strongly supportive of population structure, but population structure may be present even if the result is negative.

Given that the inclusion of baseline sequences in all clusters was enforced during the analyses, a positive result indicates that population structure was maintained following DAA treatment. Of the 80 individuals with samples that passed the analysis selection criteria, 47 (58.75%) had five or more non-overlapping windows showing evidence of structure, and 23 (28.75%) had five or more non-overlapping windows with the structure validated by both fastbaps and SAI (Figure 3, examples of structured topologies are in Supplementary Figure S5).

**Figure 3.**
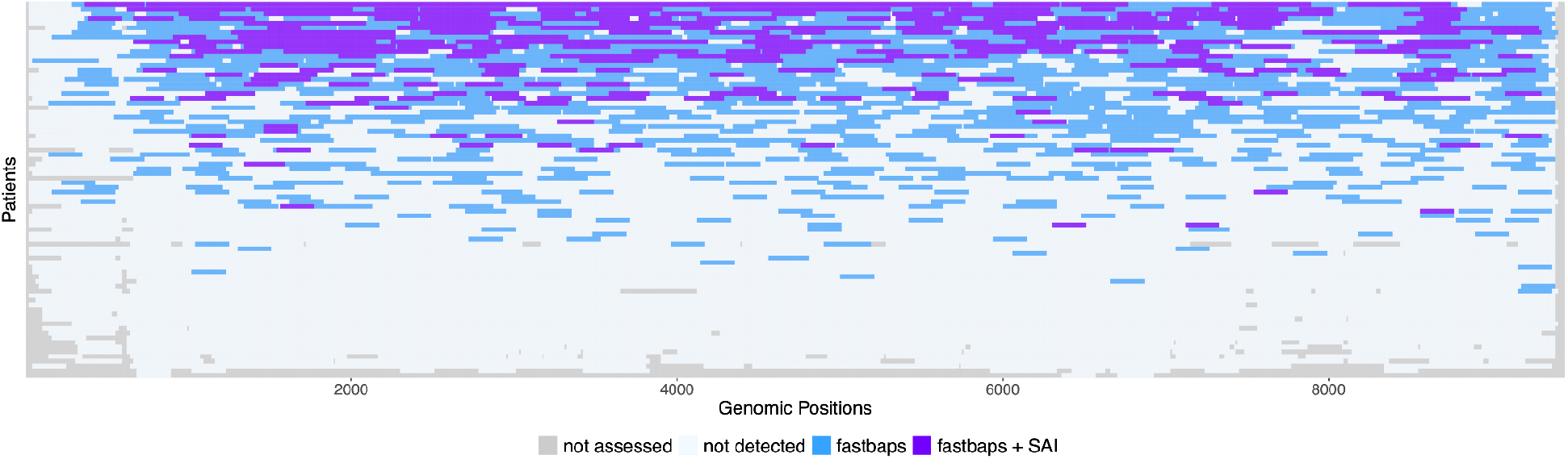
Ocean plot of windows with strong evidence for the maintenance of population structure after treatment. For sliding windows within longitudinally sampled individuals, the windows are coloured purple when evidence of population structure was maintained is strong (supported by fastbaps and SAI), are coloured blue when population structure was supported by fastbaps, are coloured light blue when the window was assessed and are coloured grey when the window was not assessed.

### Lineages that are only intermittently observed across time are less divergent

We have presented multiple lines of evidence supporting the maintenance of subpopulations after treatment in many individuals, and with the relative frequency of observed subpopulations often changing dramatically through time. Although the reconstruction of within-host haplotypes for virus populations can be challenging, the well-segregated subpopulations within HCV infected individuals appear to favour reasonably accurate haplotype reconstruction.

The reconstructed haplotype trees in our dataset showed different topologies across individuals. Some individuals had only a single lineage throughout the sampling time frame (e.g. Figure 4, patient K), whereas others had more complicated structures. For example, in some cases multiple lineages emerged from different baseline haplotypes (e.g. patient B in Figure 4), and often one or more of the lineages were observed at two sampling time points but missing from an intermediate time point (e.g. for patient B, lower lineage is missing time point PT12). We also commonly observed cases where the order of lineage branching was not chronological (e.g. Figure 4, patient L and additional examples in Supplementary Figure S6).

**Figure 4.**
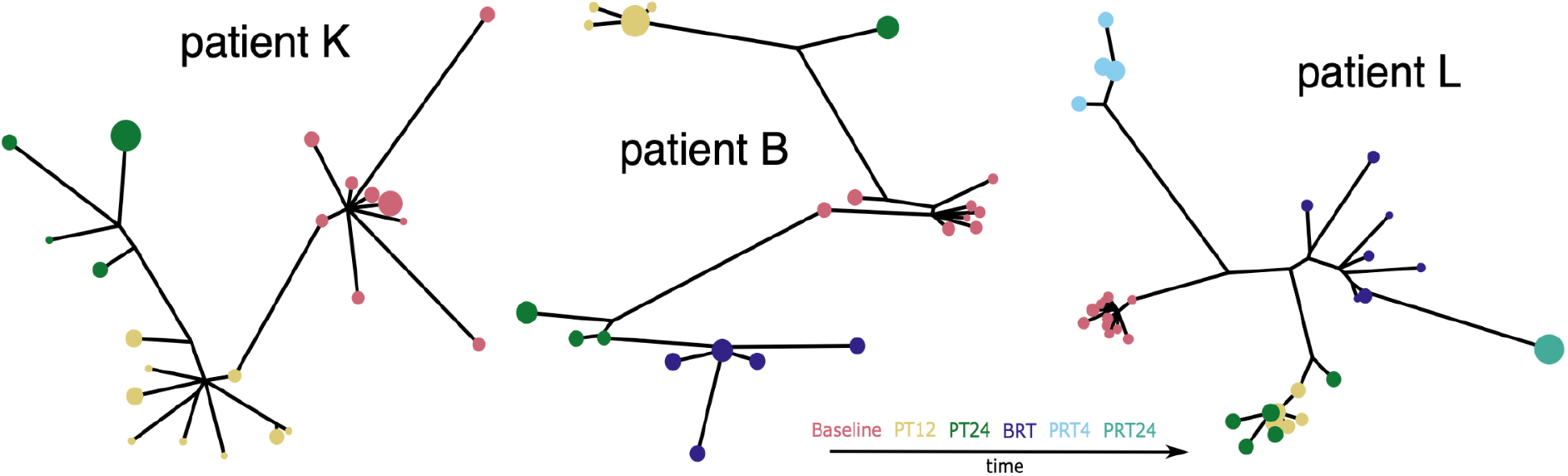
Example trees showing single subpopulation within-host (patient K), multiple subpopulations within-host (patient B), missing time point in one population (patient B: lower left lineage missing time point PT12), later sequences more similar to baseline (patient L). Baseline sequences are in red, PT12 (post treatment 12 weeks) sequences are in yellow, PT24 (post treatment 24 weeks) sequences are in green, BRT (baseline before re-treatment) sequences are in purple, PRT4 (post re-treatment 4 weeks) sequences are in light blue, PRT24 (post re-treatment 24 weeks) sequences are in light green. The size of the dots are relative haplotype frequencies.

We hypothesised that unobserved lineages may be periodically occupying different niches, and this may in turn affect their evolutionary rates. To test if there was evolutionary rate variation across the individuals, we fitted the number of synonymous mutations between haplotypes in the individual trees to two different expected distributions (Figure 5A) using a maximum likelihood framework. We used synonymous mutations as these are more likely to be neutral than nonsynonymous mutations, and therefore less likely to be influenced by selection as a direct result of treatment. A Poisson distributed number of synonymous mutations would be expected, at any given time since baseline, if all viral populations evolve at a single shared rate. Taking into account the time of sampling post baseline for all reconstructed haplotypes across all individuals in our dataset, we determined the most likely single mutation rate given the observed number of mutations on each haplotype compared to the most similar baseline haplotype. The expected distribution of mutations using the maximum likelihood rate (Poisson mean: 0.152) was significantly different from the observed distribution of synonymous mutations (Kolmogorov–Smirnov test: p<2.2×10^-16^). We next introduced additional variation into the expected distribution by fitting a negative binomial distribution. (As well as its common interpretation as the distribution of the number of failures in a given number of equal-probability Bernoulli trials, the negative binomial distribution also arises as a mixture of Poisson distributions with gamma-distributed means.) Although this expected distribution generated with the maximum likelihood parameters (shape parameter: 1.296, rate parameter: 6.931) was visually more similar to the observed distribution of synonymous mutations, the two distributions were still significantly different (Kolmogorov–Smirnov test: p=4.135×10^-6^). These comparisons demonstrated a significant overdispersion in evolutionary rates across the individuals in this study.

**Figure 5.**
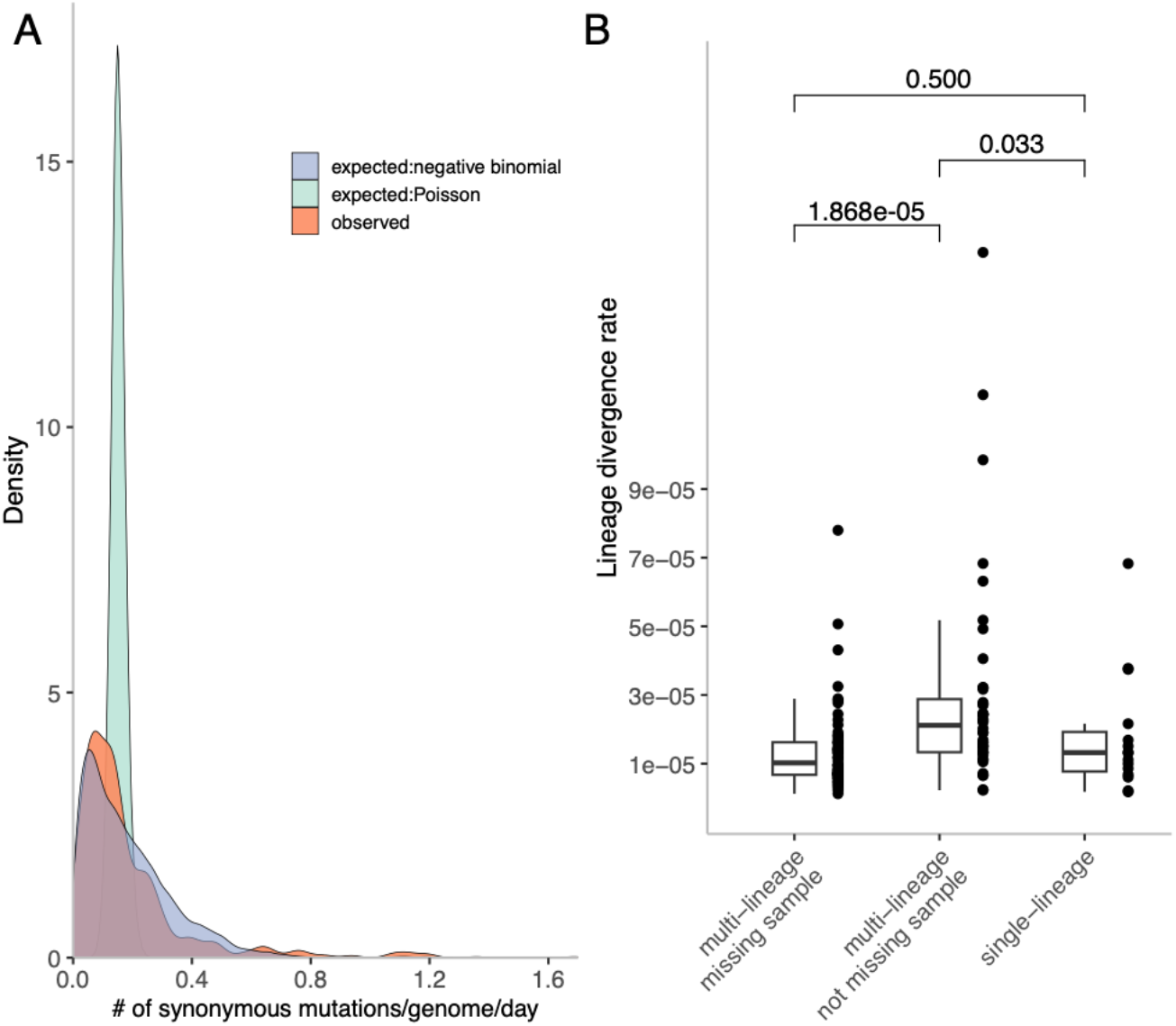
A. Distributions of the observed (red) counts of synonymous changes, a Poisson (green) and a negative binomial (blue) model generated expected counts of synonymous changes. B. Estimated rates of lineages from single lineage individuals, and multi-lineage individuals, while distinguishing between lineages where sampling time point(s) was missing or not.

To further investigate the evolutionary rate variation within individuals, we estimated evolutionary rates per within-host lineage using exhaustive model comparison of mixed-effect linear models. For individuals where a single-lineage rate model had the lowest BIC score, one rate was estimated. For individuals where a multiple-lineage rate model was the best model, we separated the lineage rates into ones that were estimated from lineages missing haplotypes from intermediate sampling timepoints and lineages that did not miss haplotypes from intermediate sampling time points. By doing so, we assumed that if the intermediate haplotype was not observed, then the subpopulation it belonged to was at low or undetectable frequency. The estimated rates for the lineages fell within the expected range for HCV neutral evolution reported by previous studies (Raghwani et al. 2019: 2.05-8.21×10^-5^ s/s/d, Gray et al. 2011: 1.11 - 1.13 ×10^-3^ s/s/yr), with lineages that had missing observations having rates significantly lower than the lineages not missing observations (Mann-Whitney U test: p = 1.868×10^-5^) (Figure 5B). These lowered divergence rates suggested that when not present in circulating blood, subpopulations may be subject to lower selection pressures, and therefore there is less opportunity for synonymous mutations to hitchhike with selected-for mutations to high frequency. Alternatively, viruses in these subpopulations might have a longer viral generation time, perhaps due to infecting different cell types. This would reduce the rate at which new mutations are introduced into the population per unit time, resulting in a lower rate of neutral evolution (Lythgoe et al. 2021). The rate estimates from single-lineage individuals were intermediate between the multi-lineage individual rates for lineages with no missing observations, and those with missing observations. This may be because the haplotypes were not well-resolved within these individuals and divergence signals from lineages of differing rates were merged into the single lineage.

## Discussion

Here, we have systematically demonstrated for the first time that HCV pre-existing within-host population structure is often maintained after DAA treatment failure. Moreover, we found considerable variation in evolutionary rates among individuals, and between different lineages within individuals, with lineages that are absent at intermediate time points tending to have lower rates of evolution. To do this we analysed whole-genome short-read virus sequences generated as part of the BOSON study, a randomised control trial to test the effectiveness of different treatment regimens of a second-generation DAA. We used a number of corroborating analyses, including tracking variant frequencies through time, scanning sliding window phylogenies across the genome, and haplotype reconstruction. This work has implications not only for our understanding of HCV biology, but also for how in the future to screen for DAA tolerance or resistance mutations before treatment and after treatment failure, which we argue should be “lineage-aware”.

Previous work has also demonstrated the presence of distinct viral lineages during HCV infections, but the question remains whether their existence poses a biological challenge for treatment efforts. Even in patients who ultimately experience viral rebound, DAA treatment induces a large reduction in the HCV viral population size, but not so large that viral diversity is not maintained; for sofosbuvir treatments at least (Foster et al. 2015). Because the resistance landscape is affected by many environmental factors such as host HLA genotype, INFL4 reactivity, DAA resistance is almost never the result of a single specific drug resistant mutation in the virus that initiated the infection (Dietz et al. 2021). The genetic resistance mechanism has proven a challenge to tease apart (Smith et al. 2021). While the cure rate of HCV infections by DAA is high, the undisrupted population structure in patients experiencing rebound after DAA treatment is concerning.

Others have characterised the mutational spectrum of HCV post DAA treatment and the development of (multi-)drug resistance from pre- to post-DAA treatment (Takeda et al. 2017; Yamashita et al. 2020; Nakamura et al. 2022). However, the role that structured populations have in the response to DAA treatment is unknown. It is possible that the presence of population structure makes the viral population more resilient to environmental disturbances caused by immunity or treatment, meaning immune- or drug-sensitive populations could be preserved within the human body, enabling them to re-emerge when immune or treatment pressures ease. One recent study showed differential persistence of RASs after treatment failure among different genotype infected individuals (Dietz et al. 2023), where high-level resistant RASs are less likely to persist compared to low- and medium-level resistant RASs. They also observed that RASs could disappear as early as 3 months after treatment failure. If it proves to be the case that population structure facilitates DAA persistence, longer treatment duration and/or treatments that target both observed and unobserved subpopulations may be advisable. If the population structure arises due to compartmentalisation, whether it be physical barriers caused by cirrhotic liver tissues or non-hepatic cells across the body, treatment strategies may need to consider the coverage of all compartments, as another study showed different RASs are present in different compartments in the liver (Sorbo et al. 2019).

Consideration of maintenance of HCV within-host population structure after treatment failure may also be critical in efforts to control the virus. In particular, we have seen time and time again that we should never be complacent over the emergence and spread of drug resistance, as witnessed by recent increases of rates of transmitted resistance to antiretrovirals in HIV infections (Pennings 2013), after decades where these rates were consistently low and considered relatively unproblematic. For HCV, there is still a large amount that is unknown, but the answers may be critical in our fight for its global eradication, which with the advent of DAAs should be our aspiration. For example, dynamic viral population structures were not seen within all the individuals we analysed, which could be because they did not exist, or because they existed but we did not observe them. Important questions remain: how widespread is within-host HCV population structure, what is the mechanism underlying the structure, why do unobserved populations have a lower rate of evolution, and what role does the viral population structure have in treatment failure and ultimately the spread of drug resistant and tolerant mutations?

It would be sensible to hypothesise that the establishment and maintenance of structured populations is complex and dependent on the interaction between host, viral, and environmental factors. Understanding the roles of these different factors could then feed into our decisions on how best to screen individuals for pre-existing virus resistance mutations, treat them, and to follow them up, so as to prevent onward transmission of resistant mutations. As a specific example, relying on the viral genome consensus at a single sampling time point for drug resistance testing risks not capturing resistant genotypes that are present but do not happen to be circulating in the blood or at sub-consensus frequency at the time of sampling. We therefore advocate for lineage awareness in evolutionary analysis of HCV infections.

Here, we analysed whole-genome short-read sequencing data. This has the advantage over previous studies looking at the within-host evolution of HCV since it covers the whole genome, rather than just the E1/E2 region that is more typically studied. However, the reliance on short Illumina reads, rather than on longer reads that are often used in this type of analysis, had its challenges because of the difficulties in establishing the physical linkage of mutations on the same genome. This meant we could not rely on the construction of phylogenies alone. We resolved this by using a sliding window approach that scanned across the whole genome, and two cross-validating statistics. This methodology ensured high specificity in the detection of population structure, but does mean there are likely many cases where population structure existed but we did not detect it with high confidence. Moreover, for part of the analysis we relied on a haplotype reconstruction tool, with the resemblance of the reconstructed haplotypes to the true haplotypes dependent on the power of the reconstruction tool and the diversity and composition of viruses within the samples. Using genuine haplotypes directly from long-read deep-sequencing data would be preferable, but while Illumina short-read sequencing is still the norm, our methods provide a roadmap for how to analyse this type of data.

Another drawback of our study is that we did not have samples pre-baseline. We therefore assumed that all subpopulations were observed at the first sampling time point. This assumption will inevitably inflate evolutionary rate estimates for lineages that were present but not observed at baseline, but which were then observed later on, since they will have appeared to have diverged a large distance from the observed baseline population (Figure 2, patient D). Future work may be able to resolve this limitation by incorporating the possibility of absent subpopulations at the first sampling time point.

In summary, we have demonstrated that within-host population structure is an important component of HCV within-host evolution in response to treatment, and we therefore advocate for lineage awareness in future evolutionary analysis of HCV. A better understanding of the within-host dynamics of HCV infections would benefit future treatment strategies, drug resistance outlook and transmission mitigation.

## Acknowledgements

The research was supported by the Wellcome Trust Core Award Grant Number 203141/Z/16/Z and 222426/Z/21/Z (to PK) with funding from the NIHR Oxford BRC. The views expressed are those of the author(s) and not necessarily those of the NHS, the NIHR or the Department of Health. KAL was supported by the Wellcome Trust and The Royal Society (107652/Z/15/Z) and the Li Ka Shing Foundation.

## Supplementary TextFile

### Method for k parameter optimisation

The *k* parameter in *phyloscanner* is used to identify collections of reads taken from the same sample which are too diverse to be the descendants of a single transmission event. This is used both to identify probable contaminant reads and to identify multiple infections. For brevity, we say this is a question of whether we are dealing with a *single event clade* or a *multiple clade event.* It works by introducing a penalty to the maximum parsimony reconstruction whereby a clade whose basal branches are sufficiently long will be reconstructed as being the result of two or more introductions to a host. When *k*=0, the reconstruction is simple maximum parsimony; increasing values of it decrease the amount of divergence along the basal branches at which the tips are split into two or more groups representing separate infection events. See the supplementary materials to Wymant et al. 2018 for full details.

The choice of a suitable *k* is not a straightforward one, particular given that *phyloscanner* is often run on hundreds of genomic windows for large numbers of samples. Branch lengths across the genome are often normalised when the package is used so that divergence between samples in different windows is on roughly the same scale. But this often leads to the branch lengths themselves being in largely arbitrary units. Even were it not so, the question of how much within-host diversity is too great to plausibly represent the result of a single infection event for any particular pathogen is a far from straightforward one. Here we outline one procedure to identify a suitable single value of the parameter for a large dataset consisting of sequences from a diverse collection of hosts.

The following procedure is intended to produce a single *k* that can be used on an entire dataset. It can be applied to the entire dataset, or if that is too large to be computationally feasible, a subset of it. The subset should be chosen to be genetically representative.

The assumption we make is that the full dataset is sufficiently diverse that if two sequences from the same host do not represent a single introduction to that individual, then they do not form a monophyletic clade in a phylogeny of all samples; the dataset’s background diversity is such that this does not happen. For small numbers of samples this may not be true, but adding sequences to make it more plausible need not be a matter of analysing more BAMs of short reads - simply adding a diverse collection of consensus sequences to the analysis will have the same effect.

In a bifurcating phylogeny, the statement that a set of *n* tips form a monophyletic clade is equivalent to the statement that the clade descended from their MRCA has 2*n*-1 internal nodes. This is also true if we are not considering tips, but the root nodes of *n* non-overlapping subclades with all child nodes pruned (such that they become tips in a subtree). If we assume that a clade is monophyletic if and only if the tips from it represent a single infection, and then this means that a set form a single event clade if and only if, for every collection of *n* nodes in the subtree descended from the MRCA of those tips, with the nodes of subtrees rooted at each of the *n* being non-overlapping, there are 2*n*-1 distinct nodes on the paths joining each of those *n* subtree MRCAs to the overall MRCA (including the overall MRCA itself).

This property can thus be used as a gold standard for testing if a clade is a single event clade. We now want to find the best value of *k* as a classifier. We can do this by, for each sample in each *phyloscanner* window (with normalised branches):

1) Prune the phylogeny so only tips from that sample remain.
2) Set *k*=0
3) Run the *phyloscanner* parsimony algorithm to split the tips into *n* groups
4) If *g*>1, record *k’=k* and the *n* MRCA nodes of the *g* groups and stop
5) If not, increase *k* by a fixed, small amount *i* (we used *i*=0.1) and go to step 3
6) Check whether there are 2*n-*1 unique nodes on the paths from the *n* group MRCA nodes to the overall MRCA node of the sample in the unpruned tree.

If there are 2*n*-1 such nodes then the sample forms a single event clade in this window according to our gold standard; if there are more than 2*n*-1 then it does not. In the former case, using values of *k* of at least *k’* result in a false positive identification of a multiple clade event, and values less than or equal to *k’-i* are true negatives. In the latter case, values of at least *k’* are true positives and values less than or equal to *k*’-*i* are false negatives. This can be used to estimate sensitivity and specificity for *k* equal to any multiple of *i*, and an optimal value for it using a ROC curve or any other means of optimising the value of a binary classifier. See figure X for an illustration.

**Figure X:**
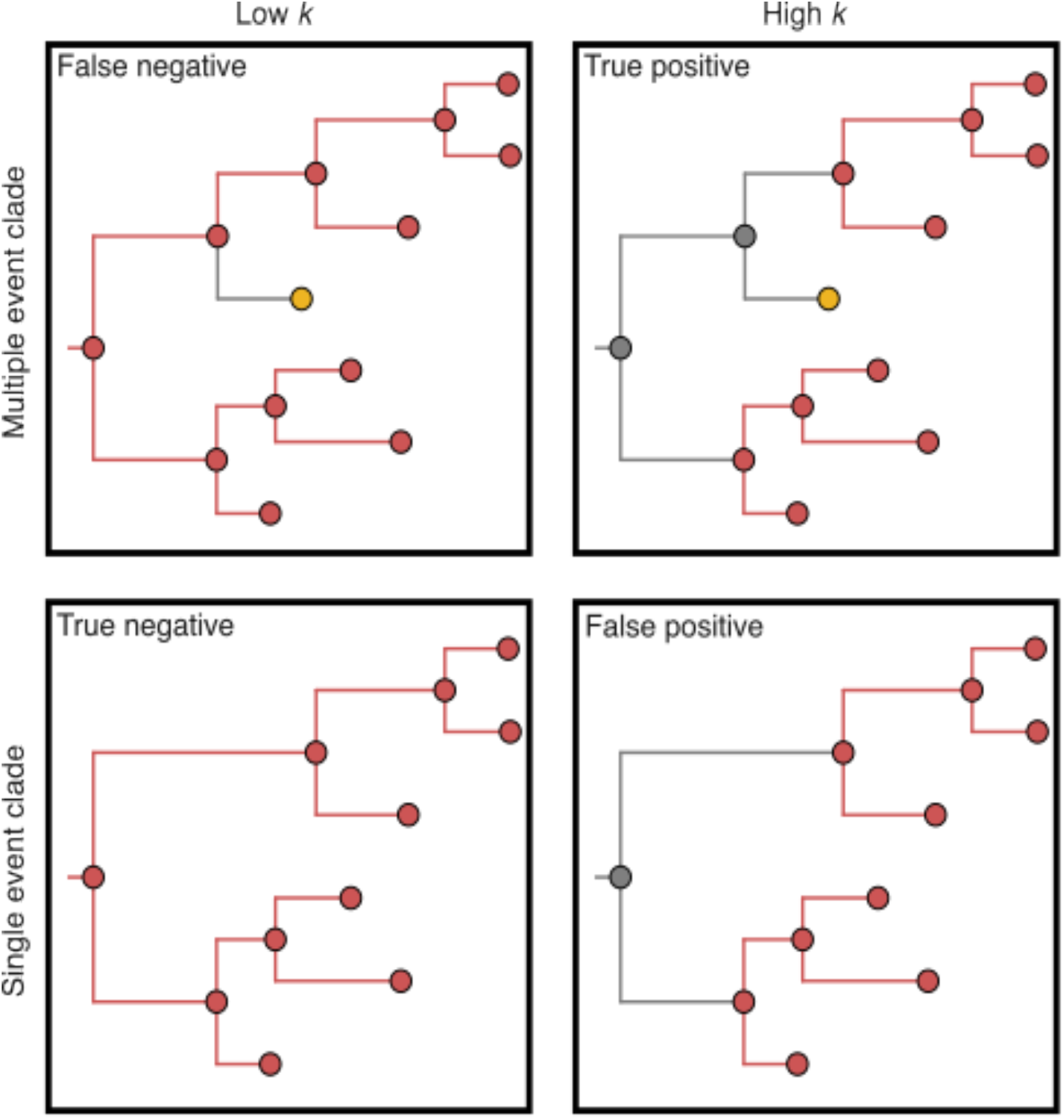
Use of *k* as a binary classifier for a phylogeny representing a multiple clade event. All red tips are derived from the same host; the yellow tip comes from a different host. It is assumed that the presence of the yellow tip indicates that this is not a single event clade, and vice versa. A *k* that is too low will miss this (top right). The yellow tip is missing from the bottom row, but a *k* that is too large (bottom right) will nevertheless declare a multiple clade event.

A naive version of this gold standard has a limitation where tips from another host are nested within the clade of interest because of precedence of the latter in the chain of transmission. In figure X this would happen if the host represented by the yellow tip was infected by the one represented by the red, possibly by way of one or more unsampled individuals. To refine the gold standard condition, we also require that, for a sample to be identified as the result of a multiple clade event, the longest branch within the subclades of the tips associated with each split is shorter than the mean of the patristic distances between the MRCAs of those splits. See figure Y for an example.

**Figure Y:**
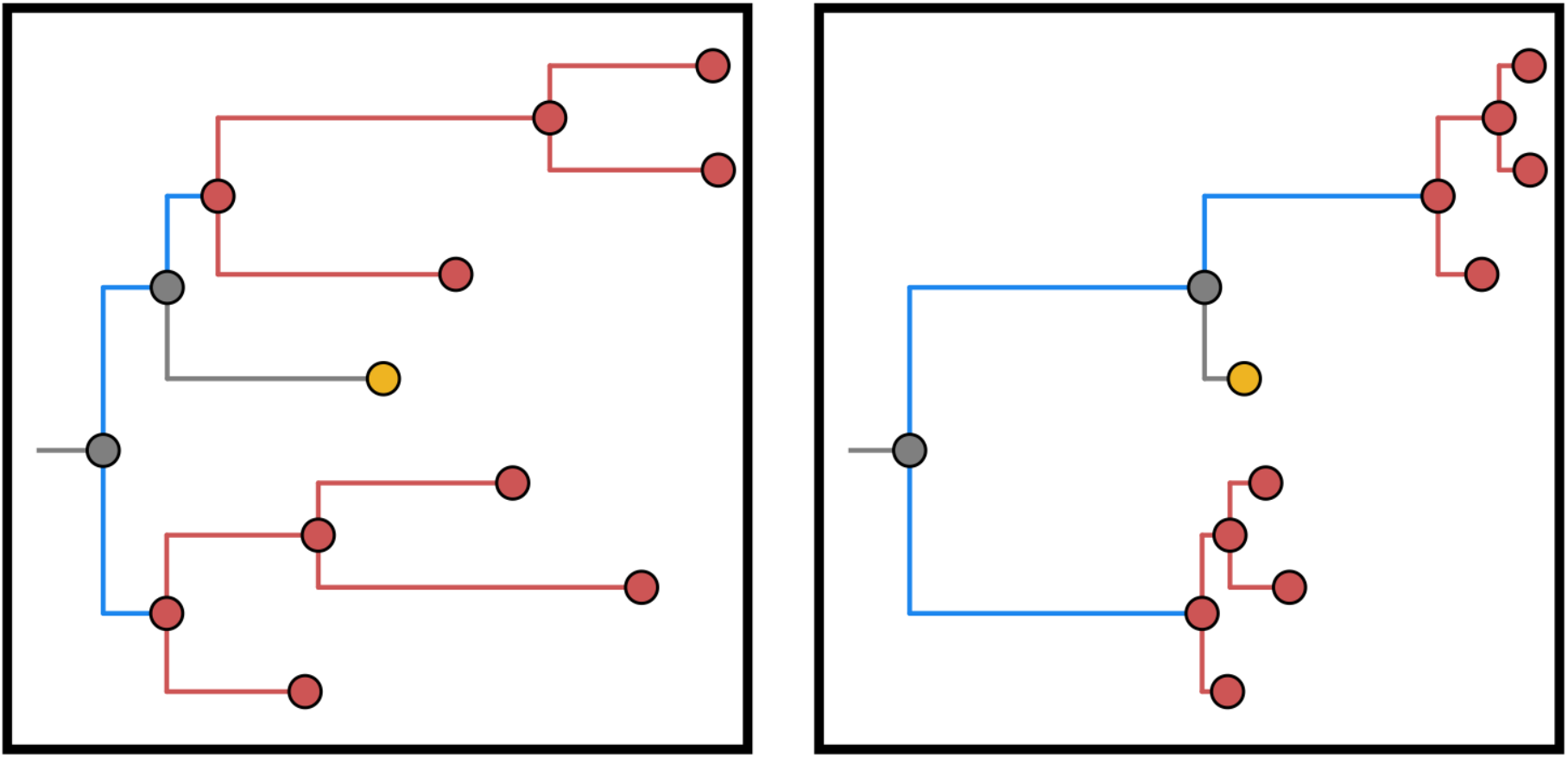
On the left, the appearance of the yellow tip nested in the diversity of the red sample is more likely to indicate the gold sample is descended from that diversity than multiple introductions to the red host occurred. The naive version of the gold standard would classify this as a multiple clade event, just as it would the right-hand tree whose basal branch lengths very much do suggest a multiple clade event. To guard against this, we also stipulate that the call of a multiple clade event requires that the longest within-clade branch (longest red branch) is longer than the patristic distance between clade MRCAs (sum of blue branches). If there are more than two such clades (not shown), the mean patristic distance between them is used.

### phyloscanner_make_trees.py command for example window 101 to 310

python phyloscanner_make_trees.py path_to_bamfiles.csv -P -A HCV_references.fasta -2 H77_NC_038882 –min-read-count 1 --x-mafft mafft --x-iqtree iqtree2 --windows 100,309

### phyloscanner_analyse_trees.R command

Rscript phyloscanner_analyse_trees.R iqtreefiles_GTR+F+R6/iqtree_ BOSON_PTs s,24 -og H77_NC_038882 -m 1E-5 -od phyloscanner_output_pbk24 -x ^([0-9]+)[0-9_A-Z]+_read_([0- 9]+)_count_([0-9]+)$ -rda -blr -ow -nr HCV_normalisation_g3a_By_Position.csv -db DuplicationData/DuplicateReadCountsProcessed_InWindow_ -tfe .treefile -pbk 24 -rwt 3 -rtt 0.01 -rcm -swt 0.5 -sdt 0.02 -amt -sat 0.33 -v 1

### cliqueSNV command

java -jar clique-snv.jar -m snv-illumina -in file.bam -log -outDir cliqueSNV_output/ -tf 0.05 -fdf extended

---

**Supplementary Figure S1.**
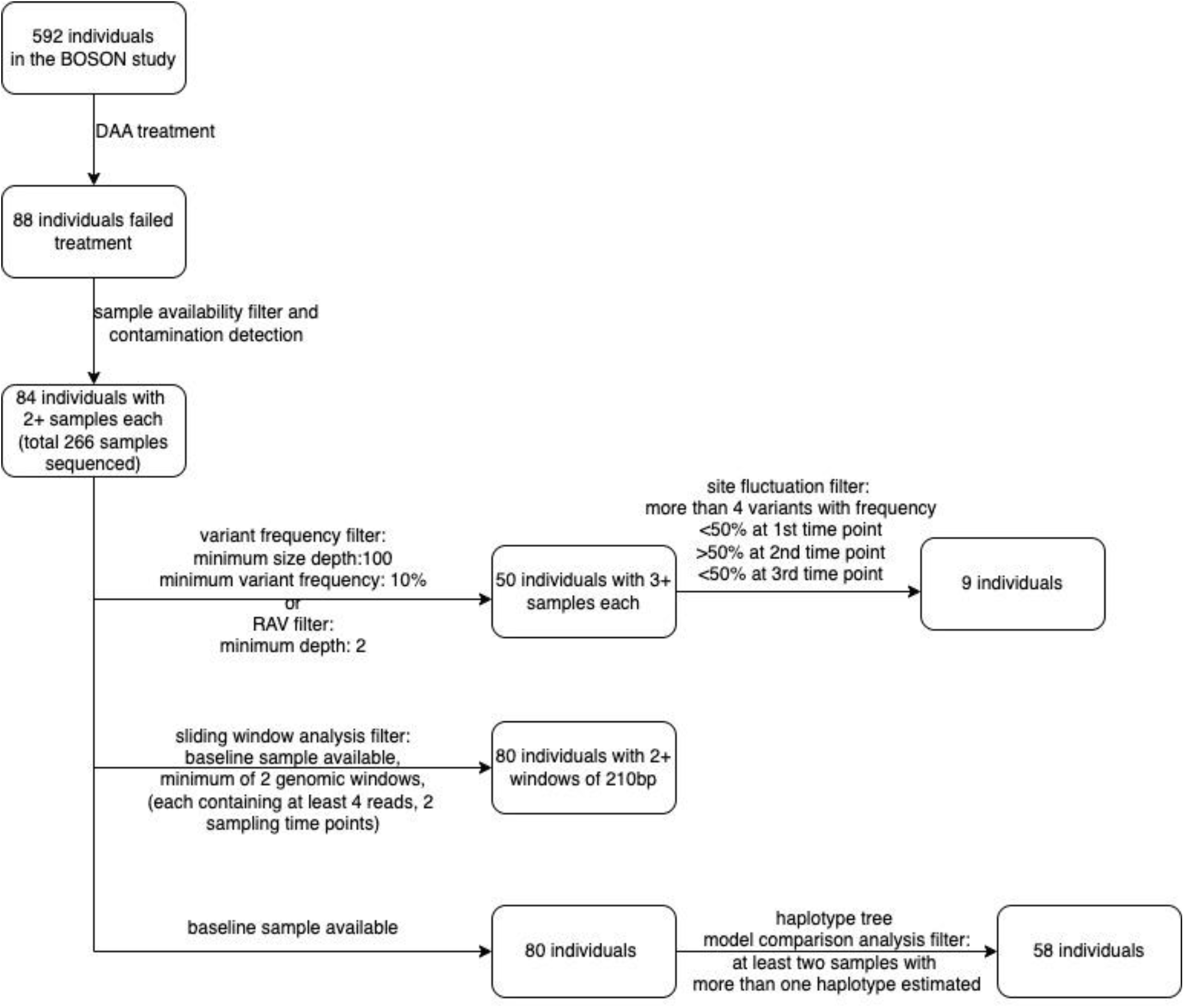
Flowchart for sample processing and quality filtering for all analyses in this study.

**Supplementary Figure S2.**
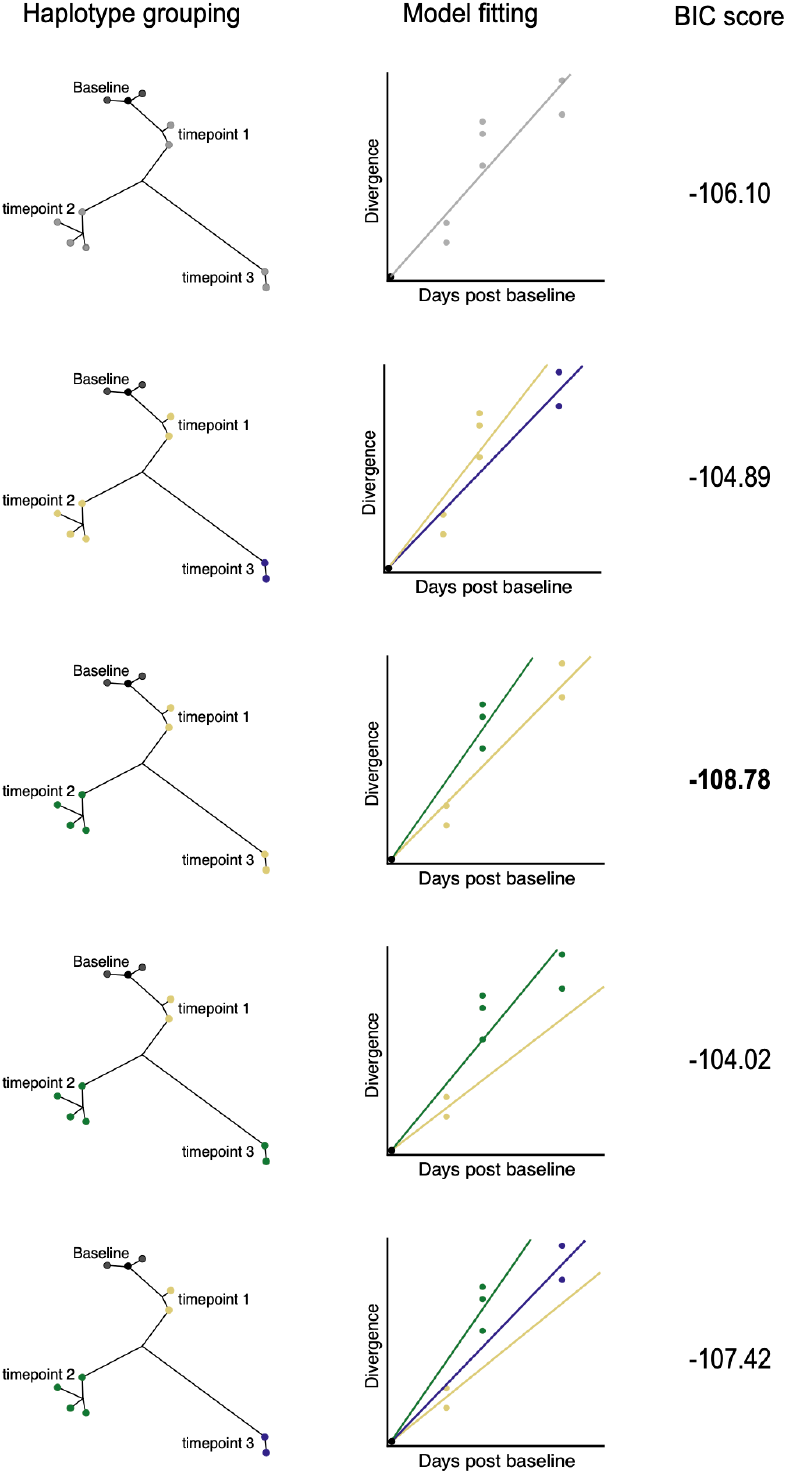
An example of the exhaustive model comparison using linear mixed-effect models for one patient’s haplotype tree. The first column lists all combinations of post Baseline haplotype grouping, as indicated by coloured tiplabels. The second column shows the fitting of the linear mixed-effect models with the corresponding groupings and the third column shows the BIC score of the model fit. The bolded BIC score of -108.78 was the lowest among all models tested and suggested that for this individual, time point 1 and 3 haplotypes belong to the same lineage, while time point 2 haplotypes belong to a different lineage.

**Supplementary Figure S3:**
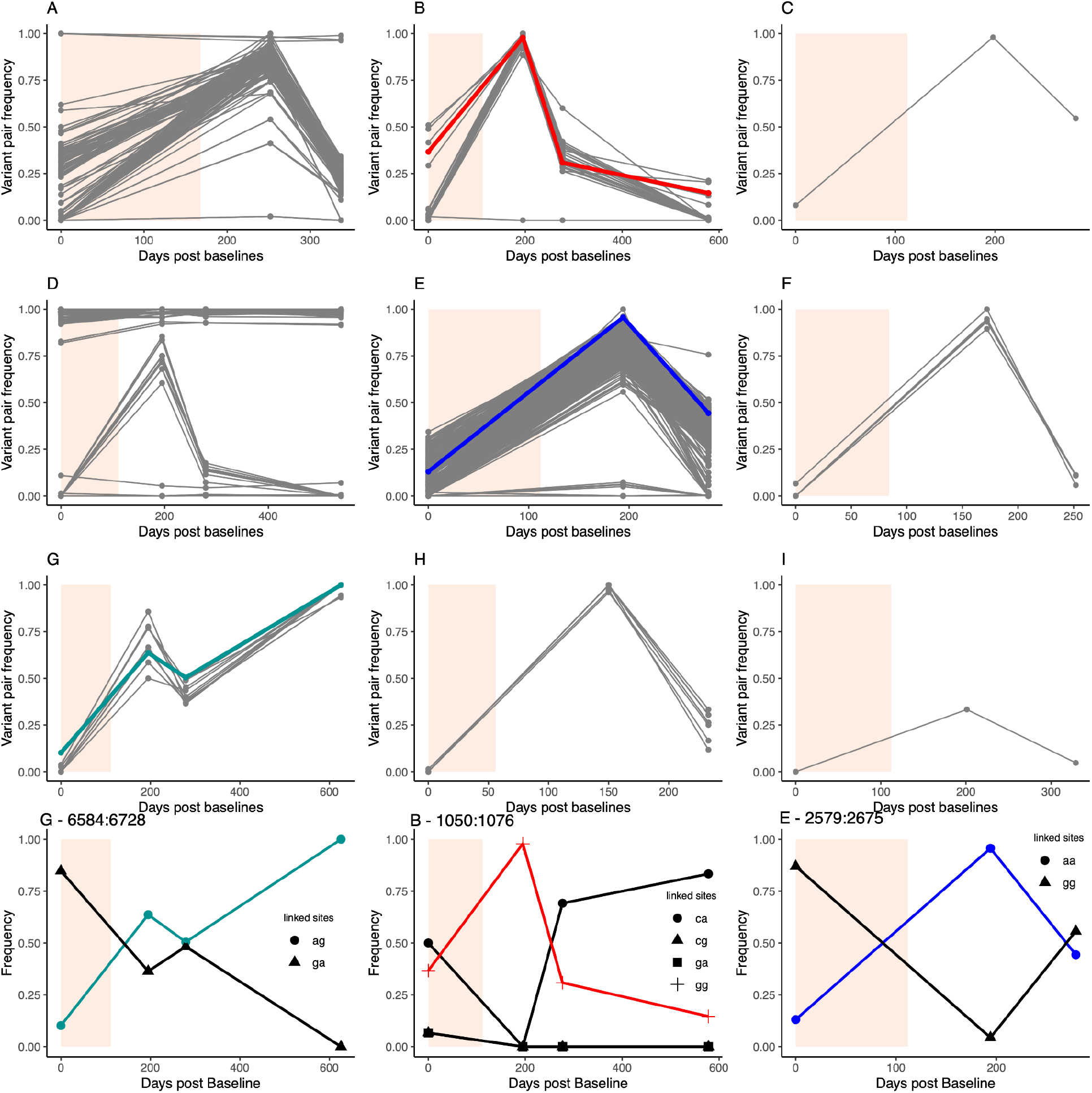
Frequency trajectories of pairs of variants that are closer than 150bp on the same sequencing fragment for patient A to I. Panel G - 6584:6728 shows the trajectories of paired variants (day 0 frequency > 5%) at genomic position 6584 and 6728 for Patient G. Panel B - 1050:1076 shows the trajectories of all paired variants (day 0 frequency > 5%) at genomic position 1050 and 1076 for Patient B. Panel E - 2579:2675 shows the trajectories of paired variants (day 0 frequency > 5%) at genomic position 2579 and 2675 for Patient E.

**Supplementary Figure S4.**
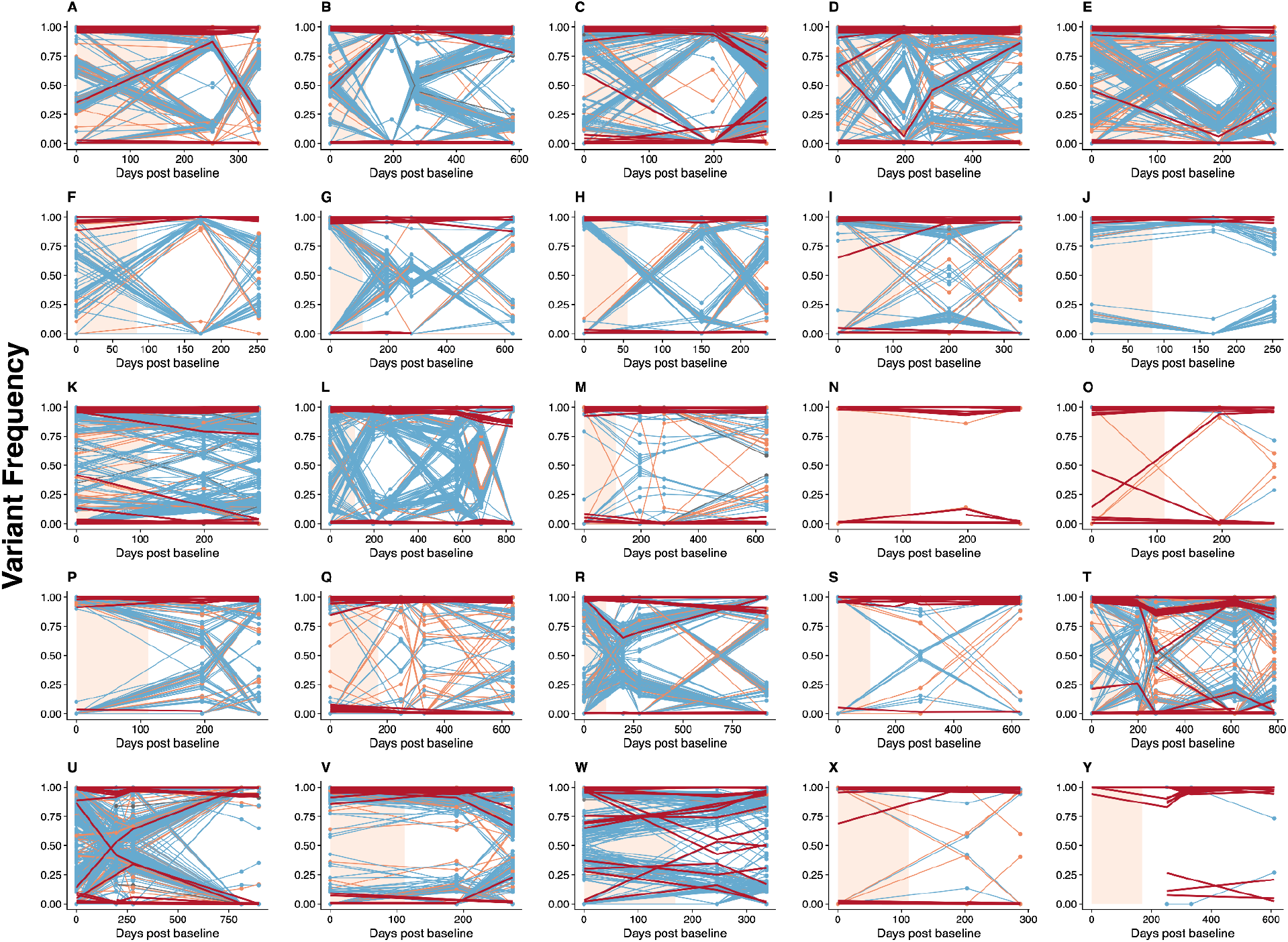
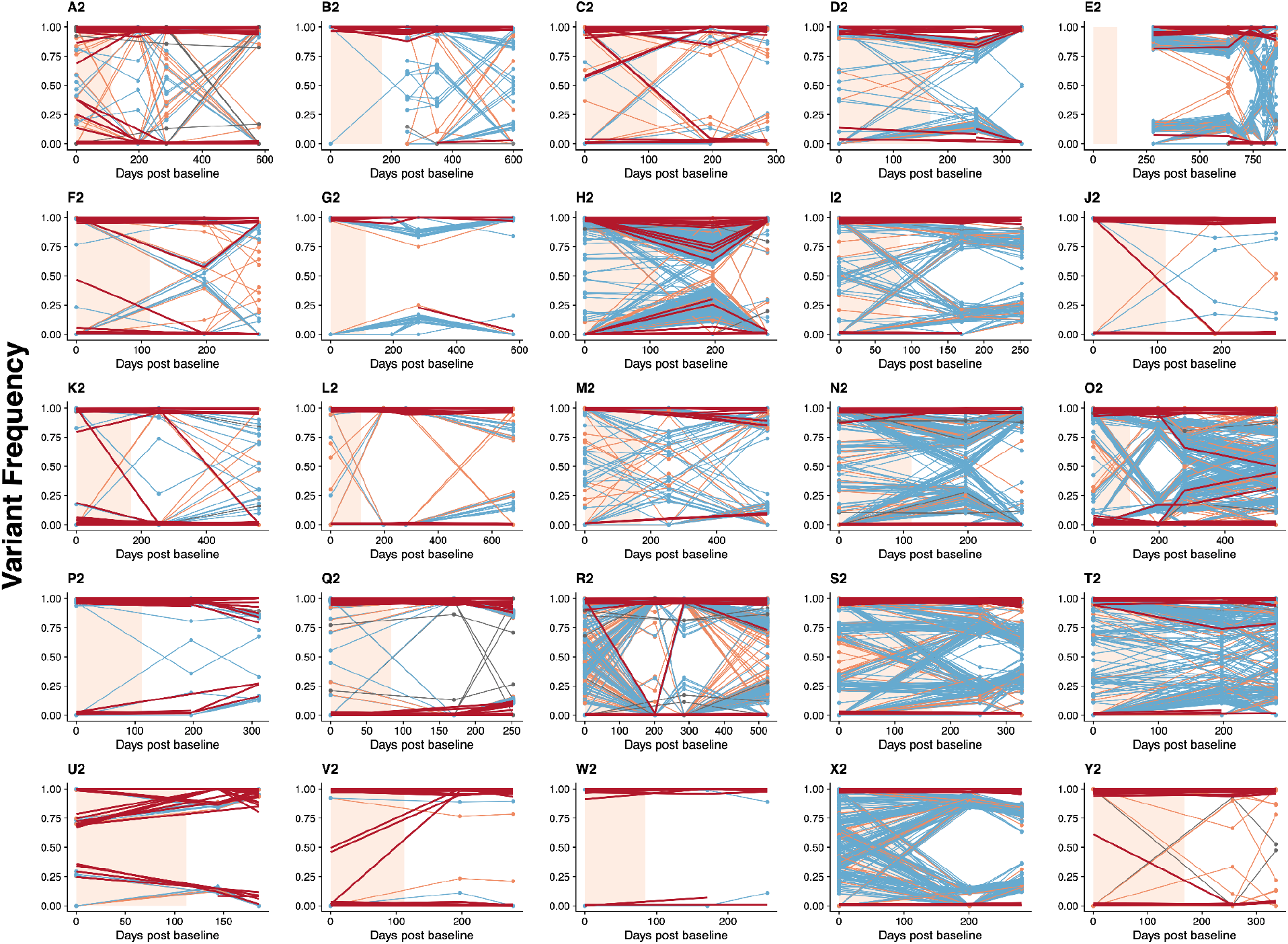
Variant frequency trajectories across sampling time points for all patients with three or more samples available. All nucleotide variants with a frequency above 10% in at least one sample were traced across all sampling time points, with synonymous changes in blue, nonsynonymous changes in orange and non-coding in grey. All relevant resistance-associated variants (RAV) trajectories are in red. The peach coloured shade indicates the duration of the DAA treatment. All variants that became fixed (frequency increased to and remained at above 90%) or were purged (frequency decreased to and remained at below 10%) after the first sampling time point were not included. Patient IDs were assigned randomly, for example, there is no relevance between patient A and patient A2.

**Supplementary Figure S5.**
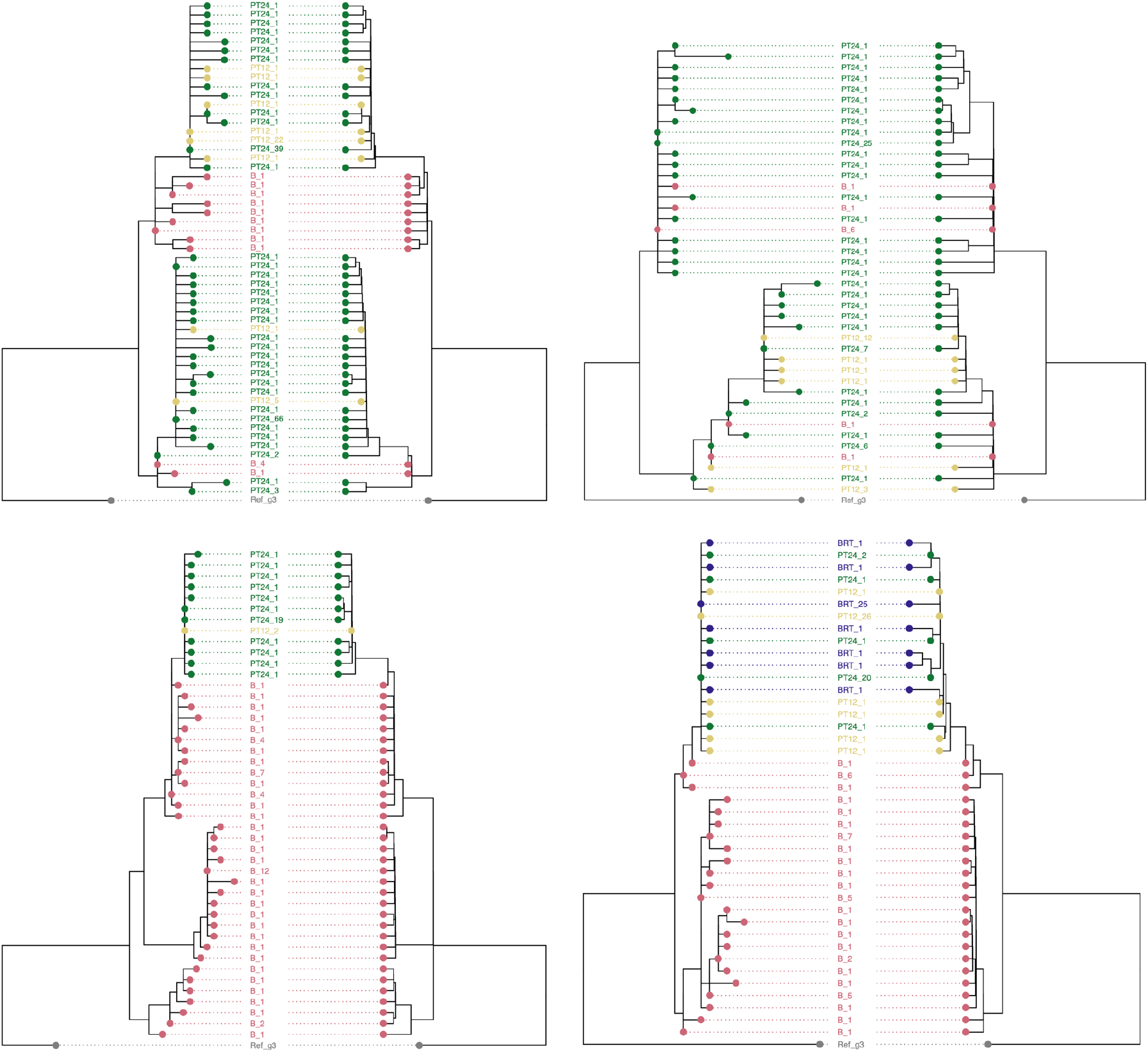
Phylogenies constructed from 210bp windows of the HCV genome sequences from four different individuals. The top two phylogenies show structure maintenance while the bottom two phylogenies do not show clear structure maintenance of the within-host viral population. The left side of each panel is the molecular phylogeny, where branch length corresponds to substitutions. The right side of each panel is the time-calibrated phylogeny, where branch length corresponds to sampling time. The tiplabels indicate the sampling time point and the number of occurrences the sequence appeared within the window. Red: Baseline (B), yellow: 12 weeks post treatment (PT12), green: 24 weeks post treatment (PT24), and blue: baseline before retreatment (BRT).

**Supplementary Figure S6.**
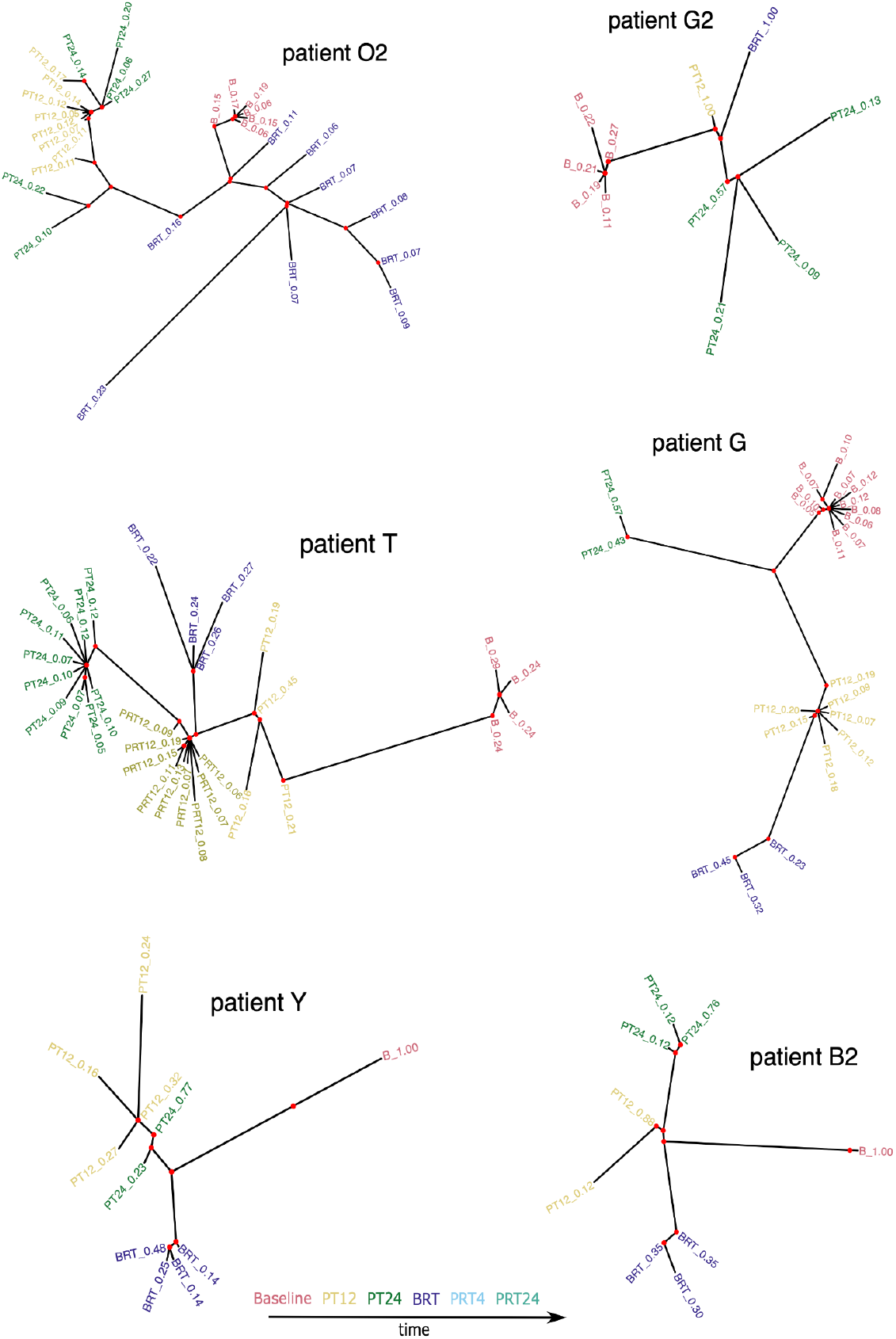
Six example patient haplotype trees where the order of lineage branching was not chronological. Baseline sequences are in red, PT12 (post treatment 12 weeks) sequences are in yellow, PT24 (post treatment 24 weeks) sequences are in green, BRT (baseline before re-treatment) sequences are in purple, PRT4 (post re-treatment 4 weeks) sequences are in light blue, PRT24 (post re-treatment 24 weeks) sequences are in light green. The tiplabels consisted of the sampling time point and the CliqueSNV-estimated frequency of the haplotype sequence from the sample. Note the haplotype frequencies from the same sample might not sum up to 1 as the minimum frequency to call a haplotype was set to 0.05.

**Supplementary Table 1.** List of 65 resistance-associated variants (RAVs) compiled from previous studies.

| Gene | Site | RAV | Relevant DAA | Publication |
| --- | --- | --- | --- | --- |
| NS5A | 28 | T | Daclatasvir | Smith et al. (2019) |
| NS5A | 30 | K | Daclatasvir | Smith et al. (2019) |
| NS5A | 31 | M | Daclatasvir | Smith et al. (2019) |
| NS5A | 32 | L | Daclatasvir | Smith et al. (2019) |
| NS5A | 54 | H | Daclatasvir | Smith et al. (2019) |
| NS5A | 58 | S | Daclatasvir | Smith et al. (2019) |
| NS5A | 62 | P | Daclatasvir | Ansari et al. (2017) |
| NS5A | 92 | K | Daclatasvir | Smith et al. (2019) |
| NS5A | 93 | N | Daclatasvir | Smith et al. (2019) |
| NS5A | 93 | H | Daclatasvir | Smith et al. (2019) |
| NS5A | 172 | E | Daclatasvir | Ansari et al. (2017) |
| NS5A | 176 | V | Daclatasvir | Ansari et al. (2017) |
| NS5A | 276 | T | Daclatasvir | Ansari et al. (2017) |
| NS5A | 294 | T | Daclatasvir | Ansari et al. (2019) |
| NS5A | 442 | S | Daclatasvir | Ansari et al. (2019) |
| NS5B | 66 | T | Sofosbuvir | Ansari et al. (2017) |
| NS5B | 90 | A | Sofosbuvir | Ansari et al. (2017) |
| NS5B | 117 | N | Sofosbuvir | Ansari et al. (2017) |
| NS5B | 120 | R | Sofosbuvir | Ansari et al. (2017) |
| NS5B | 150 | V | Sofosbuvir | Wing et al. (2019) |
| NS5B | 159 | F | Sofosbuvir | Smith et al. (2019) |
| NS5B | 180 | Q | Sofosbuvir | Ansari et al. (2017) |
| NS5B | 185 | G | Sofosbuvir | Ansari et al. (2017) |
| NS5B | 282 | T | Sofosbuvir | Smith et al. (2019) |
| NS5B | 293 | I | Sofosbuvir | Ansari et al. (2017) |
| NS5B | 316 | H | Sofosbuvir | Smith et al. (2021) |
| NS5B | 316 | N | Sofosbuvir | Smith et al. (2021) |
| NS5B | 320 | F | Sofosbuvir | Smith et al. (2021) |
| NS5B | 321 | A | Sofosbuvir | Smith et al. (2019) |
| NS5B | 401 | R | Sofosbuvir | Ansari et al. (2017) |
| NS5B | 517 | K | Sofosbuvir | Ansari et al. (2019) |
| NS5B | 571 | Y | Sofosbuvir | Ansari et al. (2019) |
| NS2 | 96 | S |  | Ansari et al. (2017) |
| NS2 | 119 | A |  | Smith et al. (2021) |
| NS2 | 131 | S |  | Ansari et al. (2019) |
| NS2 | 132 | I |  | Smith et al. (2021) |
| NS2 | 132 | V |  | Smith et al. (2021) |
| NS3 | 67 | V |  | Smith et al. (2021) |
| NS3 | 244 | R |  | Ansari et al. (2017) |
| NS3 | 256 | T |  | Ansari et al. (2017) |
| NS3 | 264 | R |  | Ansari et al. (2017) |
| NS3 | 315 | V |  | Ansari et al. (2017) |
| NS3 | 318 | T |  | Ansari et al. (2017) |
| NS3 | 354 | L |  | Ansari et al. (2017) |
| NS3 | 390 | E |  | Ansari et al. (2019) |
| NS3 | 418 | F |  | Ansari et al. (2017) |
| NS3 | 426 | I |  | Ansari et al. (2017) |
| NS3 | 559 | Y |  | Ansari et al. (2017) |
| N53 | 609 | T |  | Ansari et al. (2017) |
| NS3 | 620 | T |  | Ansari et al. (2017) |
| E2 | 61 | Y |  | Ansari et al. (2017) |
| E2 | 118 | N |  | Ansari et al. (2019) |
| E2 | 138 | A |  | Ansari et al. (2019) |
| E2 | 178 | L |  | Ansari et al. (2017) |
| E2 | 193 | E |  | Ansari et al. (2019) |
| C | 42 | P |  | Ansari et al. (2017) |
| C | 60 | E |  | Ansari et al. (2019) |
| C | 109 | P |  | Ansari et al. (2019) |
| C | 109 | S |  | Ansari et al. (2017) |
| E1 | 157 | I |  | Ansari et al. (2017) |
| E1 | 181 | T |  | Ansari et al. (2019) |
| E1 | 181 | I |  | Ansari et al. (2017) |
| NS4B | 48 | A |  | Ansari et al. (2017) |
| NS4B | 114 | S |  | Ansari et al. (2019) |
| NS4B | 162 | F |  | Ansari et al. (2017) |

